# Investigating the ATP binding pocket of CX3CL1 binding protein 2 using *in silico* approach

**DOI:** 10.1101/2023.05.24.542042

**Authors:** Rimjhim Kumari, Satinder Kaur, Rachna Hora, Prakash Chandra Mishra

## Abstract

*Plasmodium falciparum* (Pf) causes the most fatal form of malaria owing to its ability to cytoadhere in the microvasculature of various organs in the body. In addition to the Pf erythrocyte membrane protein 1 (PfEMP1) family that binds diverse host receptors, CX3CL1 binding proteins 1 and 2 (CBP1 and 2) also bind the endothelial chemokine ‘CX3CL1’ to effect cytoadhesion of parasite infected erthrocytes. CBP2 is a multifaceted protein that binds nucleic acids, Pf skeleton binding protein (PfSBP1) and ATP. ATP binding to the cytoplasmic domain of CBP2 (cCBP2) induces structural changes in the protein, and hints at its role in cell signaling. In this study, we have attempted to identify the ATP binding pocket of CBP2 using an *in silico* approach. We have also delineated the type of interactions and amino acid residues that are likely to bind ATP. As CX3CL1 binding proteins are central to parasite biology, the obtained information is likely to form the basis for inhibitor and drug design against this molecule.

## Introduction

*Plasmodium falciparum* (Pf) is responsible for causing the most fatal form of malaria in humans [1]. Asexual multiplication of this pathogen occurs inside infected RBCs (iRBCs) [2] where it exports a variety of proteins to the parasitized host cell. These include families like Pf erythrocyte membrane protein 1 (PfEMP1), Rifin, Stevor, Surfin etc. collectively termed the ‘malaria exportome’ [3]. Proteins of the exportome have diverse roles in altering iRBC rigidity and adhesiveness, their ability to uptake nutrients and escape the host immune response [4]. Cytoadherence, i.e. adhesion of parasitized erythrocytes with host endothelial cells in tissue microvasculature is chiefly mediated by the exported PfEMP1 family [5] that is reported to bind a variety of host endothelial receptors viz. CD36, ICAM-1, VCAM, CSA etc. in different tissues [6]. In 2003, Hatabu *et. al*. had identified the chemokine ‘CX3CL1’ as another receptor located on host endothelial membrane that is involved in cytoadhesion of parasitized RBCs (pRBCs) in the brain of cerebral malaria patients[7].

Constitutive expression of CX3CL1 (also called fractalkine) is seen in a variety of non-hematopoietic tissues including brain, heart, lungs and kidneys both as membrane bound and soluble isoforms [8], [9]. While the membrane-bound form of fractalkine (FKN) is an adhesion molecule, its soluble form chemoattracts monocytes and natural killer (NK) cells [10], [11]. A *P. falciparum* protein family ‘hypothetical 8’ identified by Sargeant *et al* [12] has two members that are expressed on the surface of iRBCs which act as parasite ligands involved in CX3CL1 mediated cytoadherence [13]. These are CX3CL1 binding proteins 1 (CBP1; PF3D7_0113900) and 2 (CBP2; PF3D7_1301700), which share 32% sequence identity.

CBP2 is a PEXEL positive multi-transmembrane protein that contains a nuclear localization signal as well as a cold shock DNA binding domain. Its PEXEL motif is likely to be cleaved during export leading to formation of a 2TM (two transmembrane) protein product that localizes at iRBC membranes [14]. CBP2 is also reported to co-localize with Pf skeleton binding protein (PfSBP1) at Maurer’s clefts (MCs) during the ring stage of parasite development [15]. MCs are specialized membranous structures within iRBC cytosol that are formed by the asexual parasites during their early ring stage [16], [17]. It is believed that PfSBP1 resides in Maurer’s clefts and works with other proteins like MAHRP1, Pf322, REX1, and REX2 to transport PfEMP1[18]. The cytoplasmic domain of CBP2 (cCBP2) is also known to directly bind PfSBP1, nucleic acids (DNA/RNA) and ATP [19]. Furthermore, both PfSBP1 and CBP2 have been found to be present in extracellular vesicles (EVs) produced by parasitized RBCs. EVs are responsible for transport of packaged proteins and nucleic acids to target cells and play roles in gametocyte development, inter-parasite communication and modulation of the host immunological response [20]–[24].

ATP binding of 2TM protein ‘CBP2’ is suggestive of its role as a purinergic receptor that effects cellular signaling. Binding of cCBP2 with ATP induced structural changes in the protein leading to its oligomerization [19]. ATP binding negatively impacted the interaction between cCBP2 and PfSBP1. While NMR studies identified ATP atoms H2 and H8 from adenine, and H1 of its sugar to be involved in cCBP2 binding, no information is available on the interacting residues from CBP2. Here, we have used *in silico* approach including molecular docking to identify residues from CBP2 which are likely to be engaged in interaction with ATP. Considering the importance of ATP binding sites on various proteins as valuable drug targets[25], this information may form the foundation for drug designing against CBP2.

## Materials and Methods

### Protein sequence retrieval and ATP Binding Site Prediction

The sequence of full length CBP2 was retrieved from PlasmoDB (PF3D7_1301700). ATP binding site on CBP2 was predicted by using ATPint [26]. ATPint predicts the ATP-interacting residues on a protein with Support Vector Machine (SVM) by using the sequence of the protein. The amino acid sequence (in FASTA format) of CBP2 was used as the input for prediction.

### Retrieval of structures and ATP binding site prediction

The three dimensional (3D) structure of CBP2 (PF3D7_1301700) was obtained from AlphaFold [27] via PlasmoDB [28]. Its ATP binding residues were predicted by using ATPbind [29], which is an SVM based tool that identifies ATP binding pockets based on the protein three dimensional (3D) structure.

### Molecular docking and interaction analysis

The 3D protein structure was prepared for docking by adding hydrogens and charges using UCSF Chimera 1.16 [30]. The structure of ATP was retrieved from https://pubchem.ncbi.nlm.nih.gov (PubChem CID - 5957).The 3D structure of ATP was downloaded in SDF format and further used for docking studies.

Molecular Docking was performed using Autodock via CHIMERA [30]. Both, CBP2 as receptor and ATP as ligand were prepared using CHIMERA in PDBQT format. Based on predictions from ATPint & ATPbind and information based on our report of ATP binding to recombinant cCBP2 [19], a grid box was selected to perform the docking. Others parameters for ligand and receptor were set to default [31]. After docking, the result was saved in PDBQT format. The different generated complexes were used for further analysis using LigPlot+ and Prodigy [32].

## Results and Discussion

### Sequence based ATP binding site prediction

*P. falciparum* CBP2 may be essential for parasite biology as multiple deletion attempts of its gene were unsuccessful [15], [33]. However, the genome wide *piggyBac* transposon approach showed that its gene could be mutated [34]. Keeping in mind the importance of CBP2 in a central pathogenic mechanism i.e. cytoadherence and ATP binding ability, we analysed the sequence and structure of PfCBP2 (PF3D7_1301700). Its full length sequence was downloaded from PlasmoDB and analysed using ATPint. Since the cytoplasmic domain of CBP2 spanning residues 30-160 has earlier been reported to bind ATP experimentally [19], we considered only this region for our evaluation. Arg43, Pro64, Val70, Tyr105, Arg117, Lys143, Asn 146, Lys152 and Val160 were predicted to interact with ATP.

### Structure based ATP binding site prediction

The predicted structure of full length CBP2 is available on Alpha Fold (Figure 1a). This 3D structure was verified for its structural integrity using Ramachandran plot[35] (Figure S1). All the residues of the predicted structure were found in the favoured or allowed regions. This 3D structure was used for prediction of ATP binding residues using ATPbind where structural coordinates are used as input. Here, Arg43 and Lys137 were predicted to interact with CBP2.

**Figure 1:**
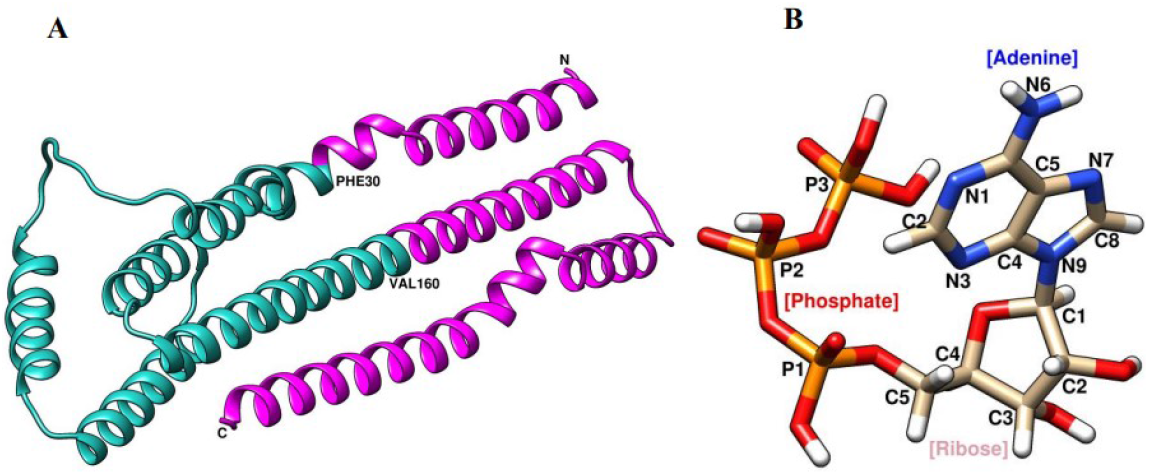
Structures of CBP2 and ATP. a) Cartoon representation of full length CBP2 obtained from Alpha fold: Cytoplasmic domain of CBP2 (cCBP2) known to bind ATP (30-160) is shown in cyan. Rest of the protein is pink. b) Structure of ATP obtained from PubChem (CID – 5957). Adenine, ribose sugar and phosphates are labeled. Carbon atoms are shown in beige, nitrogen in blue, oxygen in red, phosphorus in orange and hydrogen in white. Atoms in adenine (N1 to N9), ribose (C1 to C5) and triphosphate (P1 to P3) are separately numbered.

### Molecular docking and analysis

Since ATPint and ATPbind results showed different sites for ATP binding, we used molecular docking for understanding ATP interaction with CBP2. Molecular docking was performed by Autodock using structure of ATP ligand downloaded from PubChem (CID 5957) (Fig 1b). The CBP2 structure and ligand were prepared using Chimera for docking by generating a grid box (Fig 2 & Table S1). The grid box was selected for the docking experiment based on the result obtained from ATPint & ATPbind and previous binding report of ATP with cytoplasmic domain of CBP2 (cCBP2; residues 30-160 of full length CBP2 [19]. Nine probable complexes between CBP2 and ATP were generated by docking. PDB file of complexes generated from docking results were used for interaction studies using Ligplot+. Binding energies of the complexes were calculated using Prodigy (Table S2) [36]. The binding energies of all the complexes were found to be very similar ranging from -5.62 to -5.23 kCal/mol and docking score ranging between -7.189 to -6.138. In the complex I which had highest docking score of -7.189, ATP was seen to nestle into a deep binding pocket with adenine forming contacts with residues lying deeper in the cavity and the phosphates interacting with the more exterior amino acids (Fig. 3).

**Figure 2:**
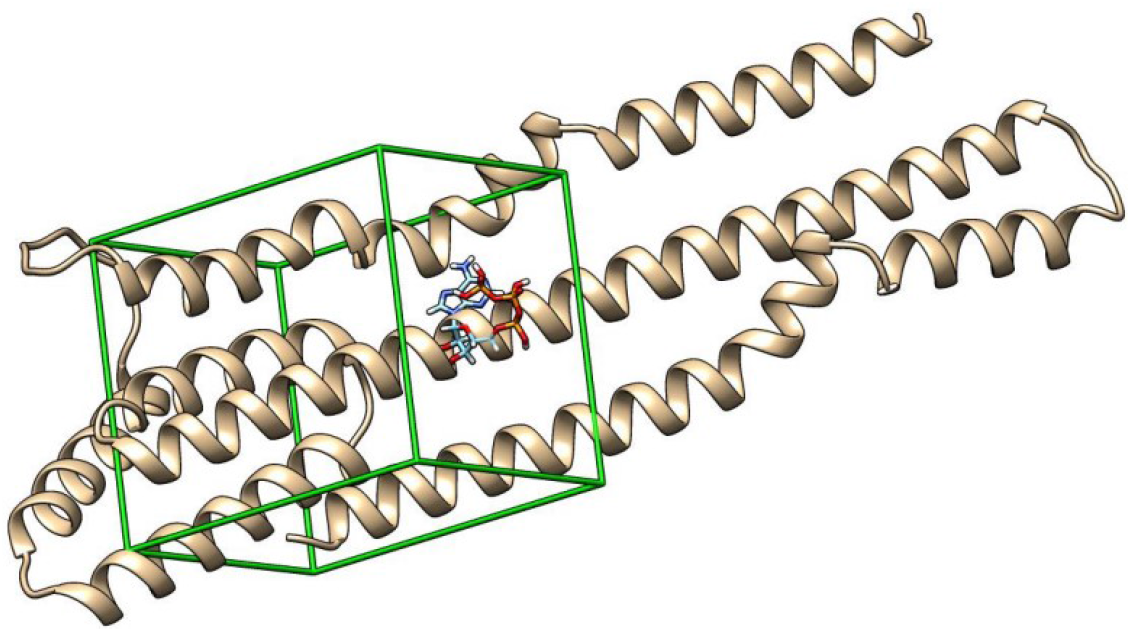
Cartoon representation of CBP2 in chimera showing the grid box and docked ATP. Inset shows selected coordinates for grid box.

**Figure 3:**
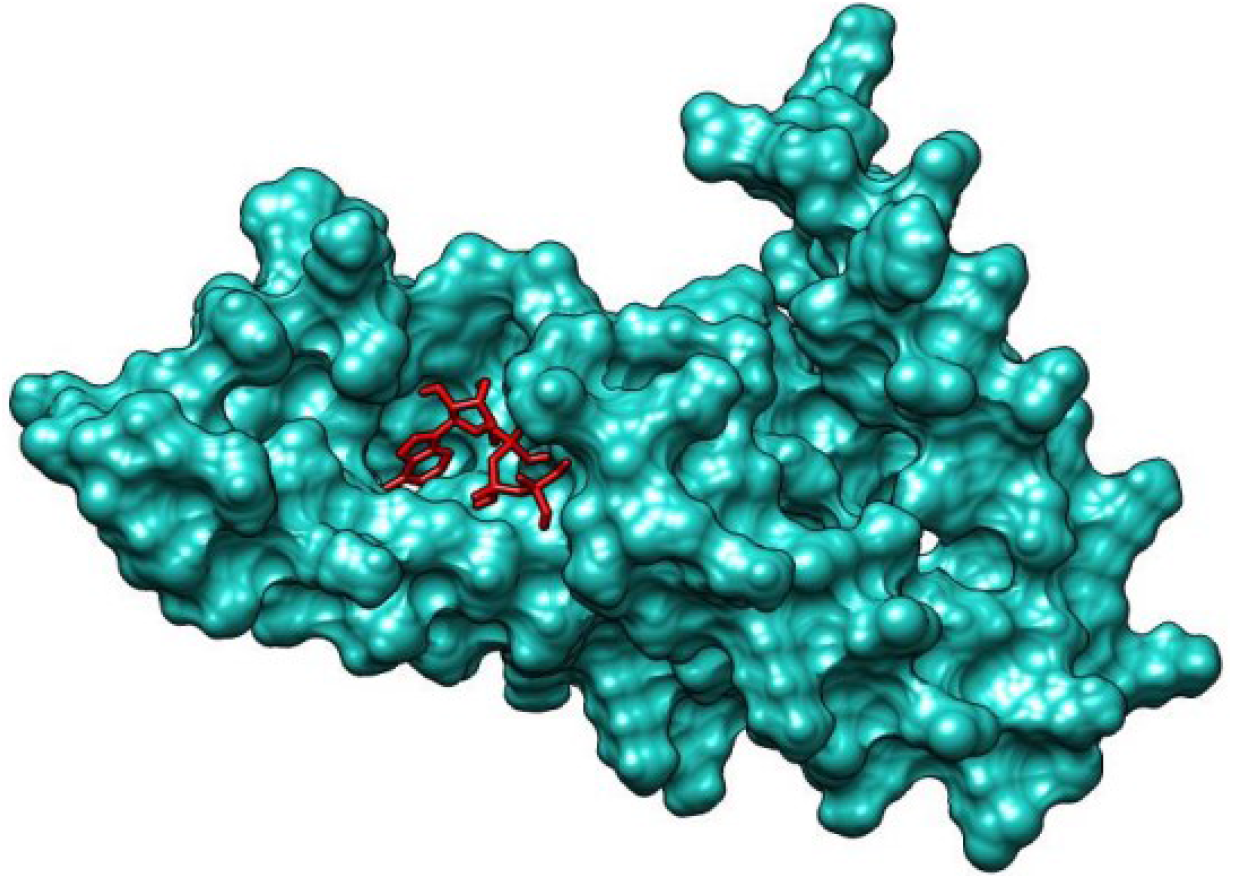
Molecular surface view of complex 1 showing CBP2 and docked ATP. Molecular surface view of CBP2 protein is shown in light blue. ATP (red sticks) are seen to bind in a deep pocket on the protein.

While complex 1 showed all the three interactions reported by NMR studies (H1 of ribose sugar; H2 & H8 of adenine from ATP) [19], rest of the other complexes showed contacts involving either one or two of these atoms. Therefore, we selected complex 1 for detailed analysis. Carbons neighboring each of the above ATP atoms were seen to be engaged in hydrophobic interactions (C1 of ribose with Arg115, C2 of adenine with Ser38 & Ile42 and C8 of adenine with Asn35) (Table 1 and Fig 4a). O atoms of different phosphate groups of ATP were seen to form hydrogen bonds. Specifically, P1 formed hydrogen bonds with Gln114, P2 with His145, Asn146 & Tyr105 and P3 with Tyr105, Lys108 & Val106 (Fig 4b). Additional observed interactions are listed in Table1.

**Table 1:** Summary of interactions in complex 1 for CBP2-ATP interaction. Table lists various hydrogen bonds and hydrophobic interactions between the two binding partners.

| Hydrogen bond |  | Hydrophobic bond |  |
| --- | --- | --- | --- |
| ATP | CBP2 | ATP | CBP2 |
| O of P1 | Gln114 | C1(Ribose) | Arg115 |
| O of P2 | His145,Asn146,Tyr105 | C2(Adenine) | Ser38,Ile42 |
| O of P3 | Val106,Lys108,<br>Tyr105 | C8(Adenine) | Asn35 |
| N3(Adenine) | Arg115 | N6(Adenine) | Ala150 |
| N6(Adenine) | Asp147 | O of P1 | Gly111,Ser110 |
| C2(Ribose) | Asp34 |  |  |

**Figure 4:**
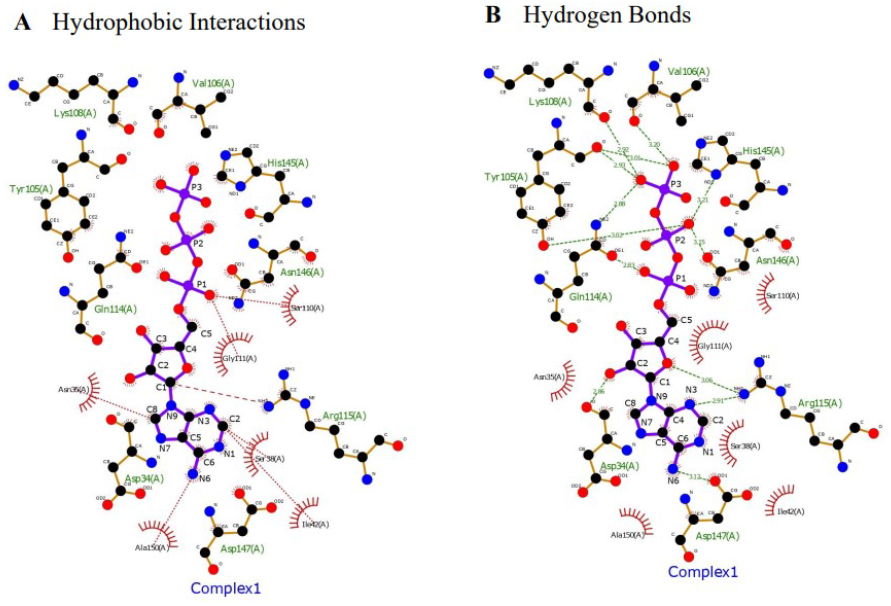
LigPlot+ analysis of complex 1. ATP is shown in the centre (ball and stick representation). Amino acid residues of CBP2 involved in interactions are shown either as ball and stick (residues forming hydrogen bonds) or as arcs (residues showing hydrophobic interactions). a) Most significant hydrophobic interactions are shown as red dotted lines. Additional residues involved in hydrophobic binding are shown with emanating rays. b) Hydrogen bonds are shown as green dotted lines. Bond length is labelled. Arg115 (shown as ball and stick) forms both hydrophobic interactionsand hydrogen bonds (a & b).

Overall, we have identified amino acid residues of CBP2 that are likely to interact with ATP by using an *in silico* approach. Our study may therefore form the basis for structure based inhibitor design against an important parasite molecule ‘CBP2’. Site directed mutagenesis of these residues can be used to confirm the importance of each of these residues in ATP binding.

## Acknowledgments

RK is receiving scholarship from DBT, Government of India. SK is a DBT-SRF. The laboratories of PCM and RH were funded by DBT, RUSA and DST, Government of India.

## Declaration

The authors have no relevant financial or non-financial interests to disclose.

The authors declare that there are no conflicts of interest. All authors read and approved the final manuscript.

## Figure legends

**Figure S1:**
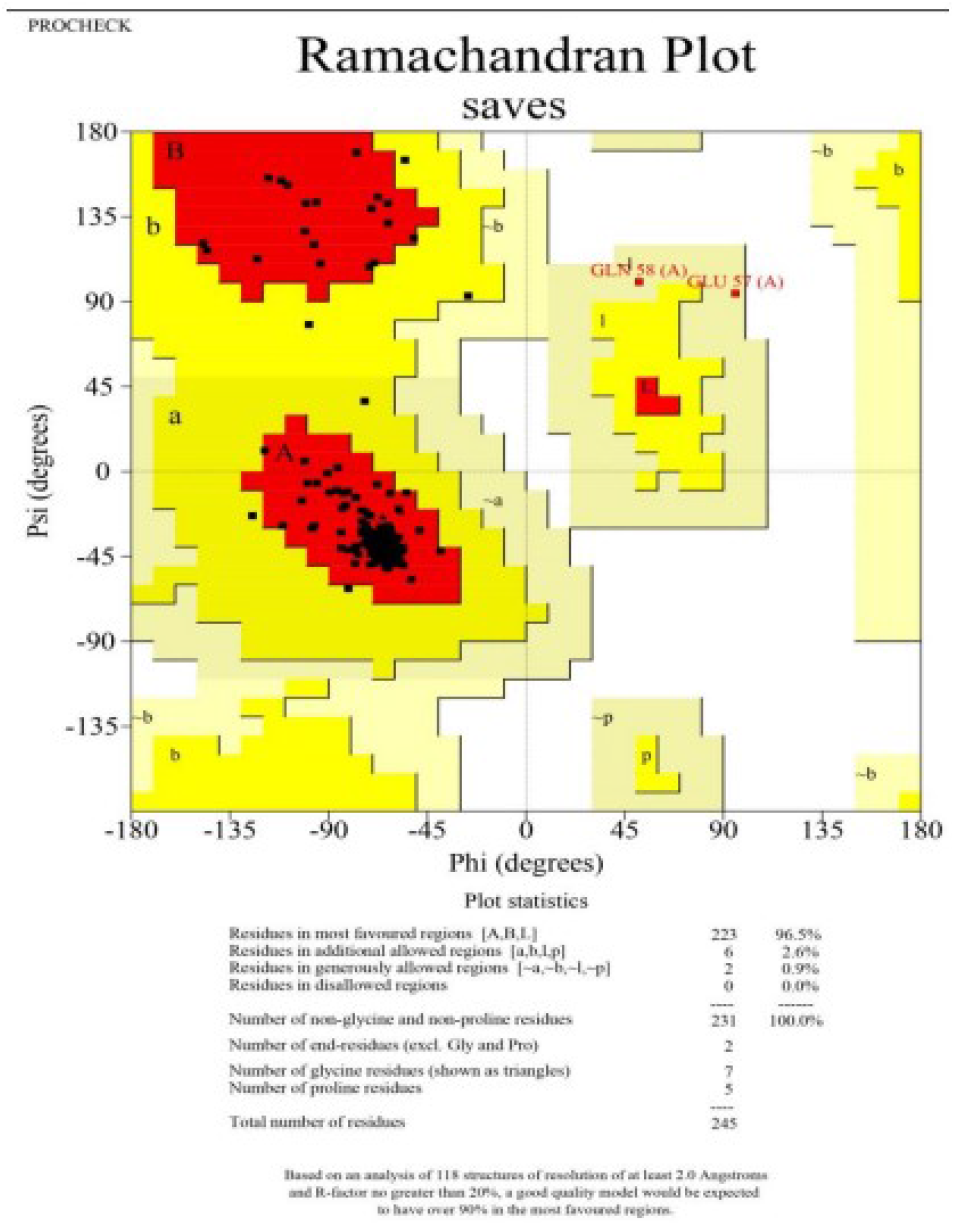
Ramachandran plot. Figure shows Ramachandran plot for predicted structure of full length CBP2 obtained from Alpha fold. All the residues are found in the allowed/ favoured regions.

**Table S1:** Grid box coordinates and dimensions. Table shows centre coordinates and size.

| Gene ID |  | X | Y | Z |
| --- | --- | --- | --- | --- |
| **PF3D7_1301700** | Center | -10.1664 | -2.87387 | 6.75774 |
|  | Size | 34.2688 | 27.7896 | 23.9081 |

**Table S2:** Summary of scores and binding energies for all complexes. Nine complexes were obtained after docking CBP2 and ATP. Scores obtained from Autodock in Chimera and binding energies (ΔG values for complexes) are listed.

| Complex Name | Score | $\Delta G$ (Kcal/mol) |
| --- | --- | --- |
| Complex 1 | -7.189 | -5.39 |
| Complex 2 | -6.924 | -5.23 |
| Complex 3 | -6.841 | -5.39 |
| Complex 4 | -6.548 | -5.49 |
| Complex 5 | -6.483 | -5.62 |
| Complex 6 | -6.466 | -5.45 |
| Complex 7 | -6.45 | -5.51 |
| Complex 8 | -6.375 | -5.47 |
| Complex 9 | -6.138 | -5.31 |

## Notes

### Competing Interest Statement

The authors have declared no competing interest.

